# Assessing the Use of Antiviral Treatment to Control Influenza

**DOI:** 10.1101/001537

**Authors:** Sarah Kramer, Shweta Bansal

**Affiliations:** Department of Biology, Georgetown University, Washington, DC 20057, USA; Fogarty International Center, National Institutes of Health, Bethesda, MD 20893, USA

**Keywords:** influenza, A(H1N1)pdm09, epidemiology, public health, antiviral, treatment, vaccine, susceptibility, infectivity, pandemic

## Abstract

Vaccines are the cornerstone of influenza control policy, but can suffer from several drawbacks. Seasonal influenza vaccines are prone to production problems and low efficacies, while pandemic vaccines are unlikely to be available in time to slow a rapidly spreading global outbreak. Antiviral therapy was found to be beneficial during the influenza A(H1N1)pdm09 pandemic even with limited use; however, antiviral use has decreased further since then. We seek to determine the role antiviral therapy can play in pandemic and seasonal influenza control on the population level, and to find optimized strategies for more efficient use of treatment. Using an age-structured contact network model for an urban population, we find that while a conservative antiviral therapy strategy cannot replace a robust influenza vaccine, it can play a role in reducing attack rates and eliminating outbreaks.

## 1 Introduction

Influenza causes yearly epidemics, and has been responsible for four pandemics during the last 100 years [1, 2]. Infection leads to high levels of severe complications and death in both young children and the elderly [2, 3]. Vaccines are widely accepted as the best tool to combat influenza [2]; however, current vaccines exhibit several shortcomings. A new vaccine must be developed each year, and circulating strains must be predicted months before an epidemic occurs. In the case of a pandemic, an effective vaccine is unlikely to be available until several months after the pandemic begins [4]. A recent meta-analysis by Osterholm et al. revealed the influenza vaccine efficacy found in ten randomized controlled trials to be 59% on average, and furthermore found little or no evidence of vaccine efficacy among those under the age of 18 or over the age of 64 [5, 6]. While this finding by no means shatters our vaccine-led influenza control efforts, it reminds us of the need to continue to develop better influenza vaccines and therapeutics, as well as more efficient strategies to distribute these interventions.

Antiviral drugs, such as the neuraminidase-inhibitors oseltamivir and zanamivir, may be attractive alternatives to current vaccines. Unlike influenza vaccines, antiviral drugs are not strain-specific [7]. When used to treat infected individuals within 48 hours of symptom onset, antivirals reduce the probability that an infected individual will transmit influenza to his or her contacts [8, 9, 10, 11]. In this study, we aim for a systematic understanding of the population-level impact of antiviral usage, and seek to answer the following questions. 1) For seasonal influenza, can antivirals replace vaccination, especially in the case of poor vaccine match or a shortage in vaccine supply? 2) In the case of a pandemic, when faced with no vaccine for several months, what impact will the use of antivirals have? 3) Are there focused uses of antivirals that optimize their impact? Without a greater understanding of the potential transmission-reducing effects of antiviral drugs, we cannot confidently declare vaccination to be the most effective influenza control strategy available.

Previous modeling studies assessing the population-level impact of antivirals have helped establish the potential of antiviral treatment as an influenza control strategy, but most of these studies reveal greatly varying results. Carrat et al. and Ferguson et al. find that treating 63% or 45% of clinically ill individuals with antivirals leads to only a 7% or 15% reduction in pandemic size, respectively [12, 13], while a study by Pepin et al. finds that 40% coverage of infected individuals can reduce seasonal epidemic transmission by 30% [14]. A study by Longini et al. has also considered the impact of prophylactic treatment of close contacts of infected individuals, and finds that prophylaxis of 80% of all exposed individuals is nearly as effective as vaccinating 80% of the population [15]. While this study shows antiviral treatment to be an effective control measure, it relies on specific contact prophylaxis based on unrealistically-intensive contact tracing. Finally, a study by Black et al. published earlier this year uses mathematical modeling to better understand why antivirals were ineffective at containing the A(H1N1)pdm09 pandemic, and finds that early treatment and prophylaxis of individuals in infected households is crucial if influenza transmission and pandemic doubling time is to be significantly reduced [16]. This study is closest in nature to our own, aiming for a quantitative assessment of the use of antivirals in a simple epidemiological model with social structure. Our study differs from the work of Black et al. because we aim not to explain the lack of population-level impact of antivirals during the A(H1N1)pdm09 pandemic, but rather to consider the scenarios under which antiviral use would be viable in future pandemics as well as seasonal epidemics. Using a network model where individual hosts are “nodes”, and interactions (i.e. contacts) that may allow influenza transmission are “edges” (details in Methods below and Figure 1), we simulate SIR epidemics to assess the impact of antiviral treatment strategies on influenza control. We also develop a parallel intervention model of trivalent inactivated vaccine (TIV) use, to consider the impact of antivirals as judged against the well-understood and widely-accepted case of vaccination.

**Figure 1:**
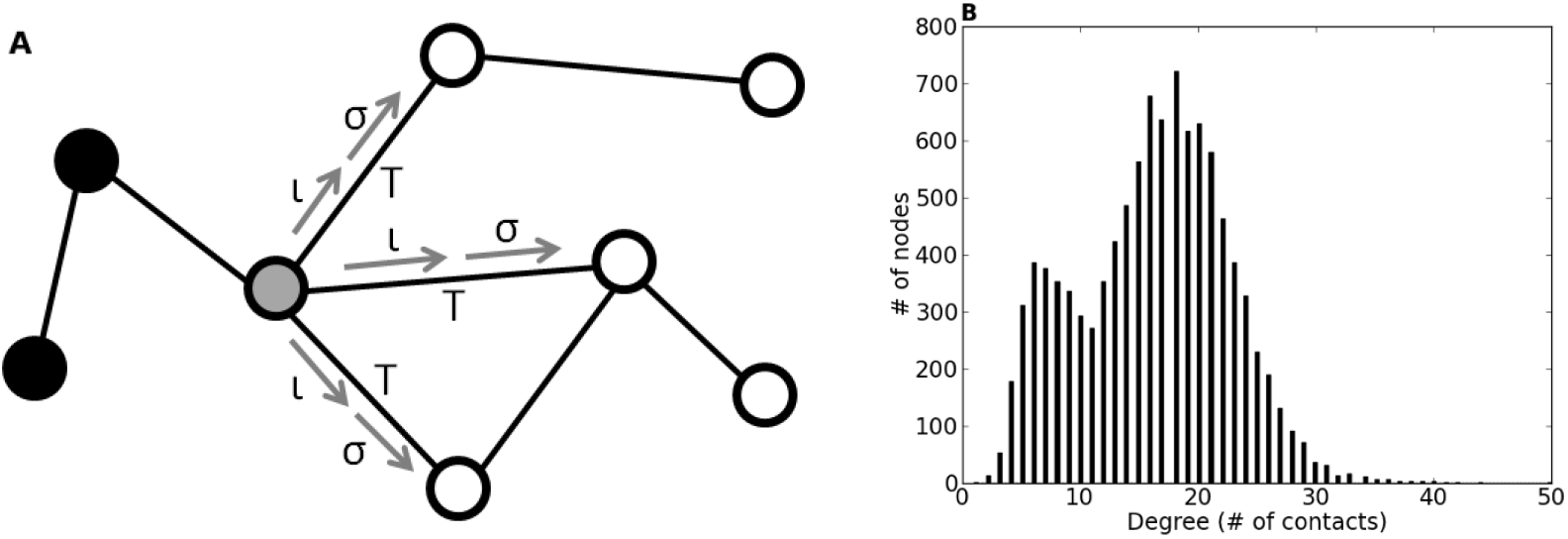
Urban contact network schematic and connectivity profile: (A) Simple example of a contact network model, where circles (i.e. *nodes*) represent individuals and lines connecting them (i.e. *edges*) represent contacts over which influenza can spread. Black nodes are recovered, gray are infected, white are susceptible. Infected nodes infect susceptible contacts with probability *T* = *ισ* (where, *T* is the transmissibility, *ι* is infectivity, *σ* is susceptibility). (B) The frequency distribution of number of contacts per individual, or *degree* in the urban contact network model. The network contains 10,304 individuals with an average degree of 16.11.

## 2 Results

### 2.1 Random Vaccination and Antiviral Treatment

To assess the potential of antiviral treatment in reducing epidemic and pandemic attack rates, we first compare the impact of random allocation of either vaccination or antiviral treatment. Vaccination is implemented at the coverage levels specified (Table 1), and antiviral treatment is implemented at a range of coverage levels (Figure 2). Antiviral coverage is reported as a proportion of the infected population size (e.g. 20%, 40% etc.) unless otherwise specified.

**Figure 2:**
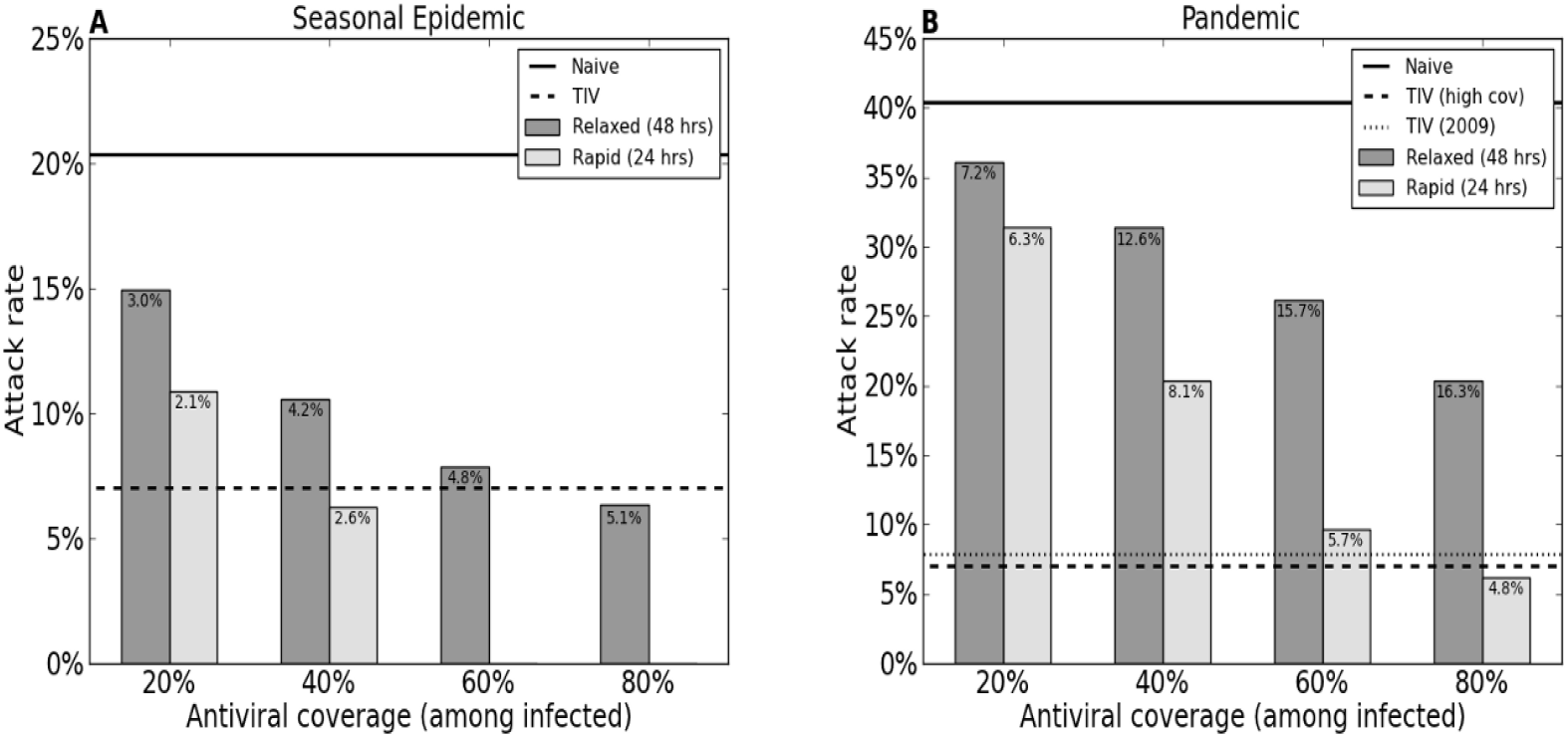
Attack rates when antivirals are allocated randomly during (A) seasonal epidemics and (B) pandemics. Horizontal lines show attack rates in populations using no control strategies (“naive”; solid lines) and vaccination (dotted lines; in (B), both model pandemic vaccines are shown). Both relaxed (dark gray) and rapid (light gray) scenarios are displayed. Percent of infected individuals treated is shown along x-axis, and percent of total population treated is shown within the bars. Results are only shown for those simulations in which at least 5% of the population was infected. Error bars are not shown, as standard errors for attack rates were all below 0.008.

TIV = trivalent inactivated vaccine

**Table 1.** Efficacy and coverage values for each influenza control scenario used (values for efficacy and realistic vaccine coverage are weighted averages of values for all age gruops)

| Control scenario | Efficacy | Coverage |  |
| --- | --- | --- | --- |
|  |  | Random | Realistic |
| Seasonal vaccine | 64.5% | 20% | 24.7% |
| 2009 vaccine | 79.5% | 30% | 29.1% |
| High coverage vaccine | 64.5% | 40% | 45.4% |
| Relaxed antiviral treatment | 20% | Varies | 30% |
| Rapid antiviral treatment | 40% | Varies | 30% |

When the relaxed antiviral treatment strategy is employed against seasonal influenza, 80% coverage of infected individuals is required to reduce epidemic attack rates by the same amount as random vaccination. When the rapid strategy is used, 40% coverage is required, and no large-scale epidemics emerge when coverage is 60% (Figure 2a). In the case of a pandemic, the relaxed strategy does not reach the population-level effectiveness of either pandemic vaccine at any coverage level tested, and the rapid strategy is more effective than both vaccines only at 80% coverage level (Figure 2b).

Figure 2 also displays the percent of the total population treated in each scenario. As coverage of infected individuals increases, these values first increase, then decrease as fewer individuals become infected, suggesting that coverage among infected individuals, and not among the population as a whole, is important for population-level antiviral effectiveness. Furthermore, the rapid strategy consistently outperforms the relaxed strategy, even though fewer members of the overall population are treated, indicating the importance of initiating treatment soon after symptom onset.

### 2.2 The Limits of Vaccination

We next consider the impact of vaccine efficacy on influenza control to determine the minimum overall vaccine efficacy beyond which antiviral use is more effective at reducing attack rate. Here, we assume 30% antiviral coverage (similar to coverage observed during the A(H1N1)pdm09 pandemic [17, 11, 14]) and realistic age-based vaccine coverage levels (Table S1). We choose realistic vaccine coverage levels to better illustrate the actual population-level impact of vaccines, rather than their impact were they to be distributed ideally. In Figure 3, we find that vaccines begin to reduce seasonal epidemic attack rates by more than random, realistic antiviral treatment at vaccine efficacies between about 25% and 50%, depending on the timing of antiviral treatment. During a pandemic, vaccines begin to outperform antivirals at efficacies between approximately 10% and 35%, depending on treatment timing.

**Figure 3:**
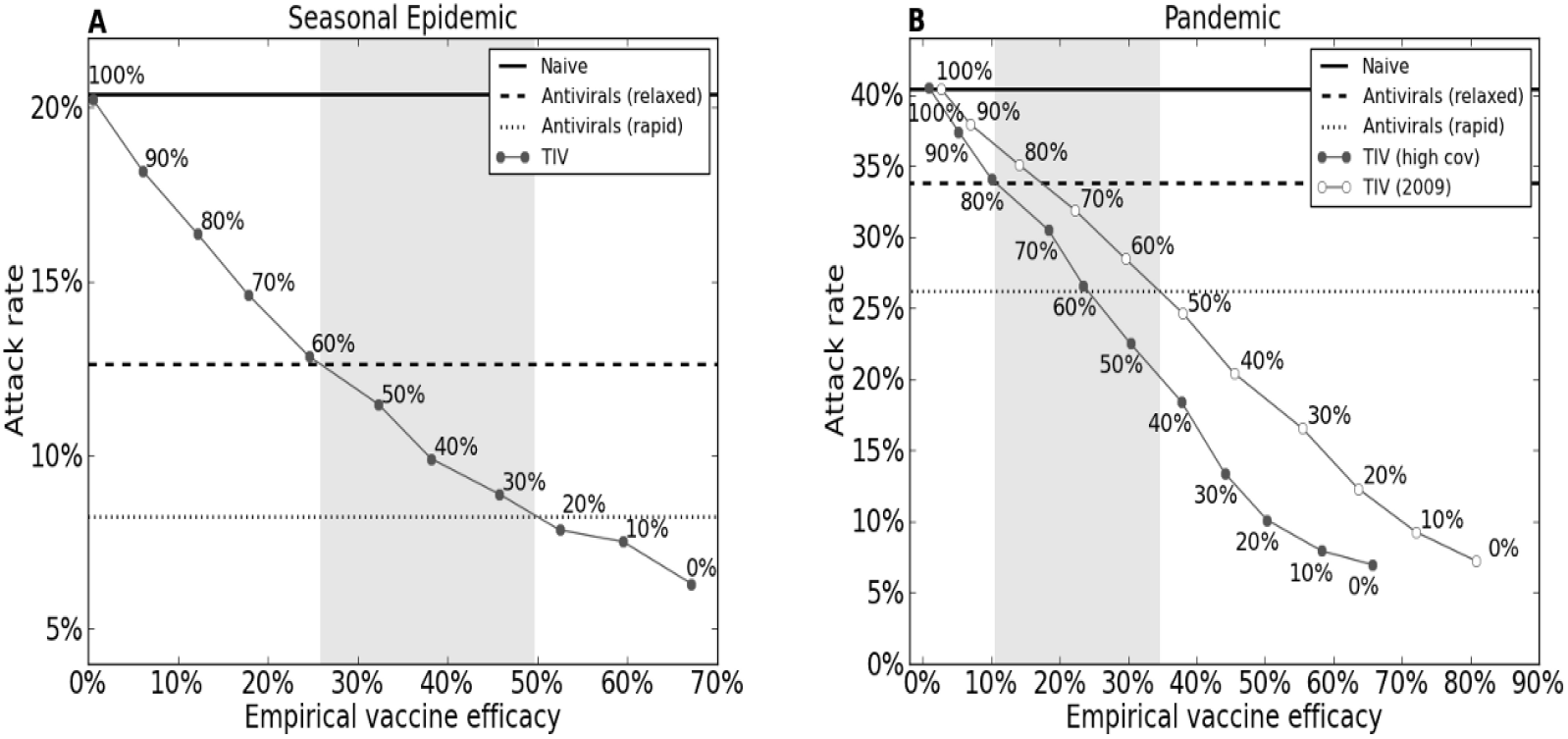
Attack rates for (A) seasonal and (B) pandemic influenza when vaccines of varying efficacy are employed at realistic, age-based coverage levels. Horizontal lines show mean attack rates for naive (solid line) populations and populations using antiviral treatment at realistic 30% coverage (dotted lines; both scenarios shown). Sloped lines show mean attack rates for populations using vaccine; both pandemic vaccines are shown in (B). Vaccine efficacies displayed on x-axis, percent reduction in individual-level vaccine efficacy for all age groups beside each data point. Again, results are only shown for simulations in which at least 5% of the population was infected. Standard errors were consistently below 0.003.

### 2.3 Focused Antiviral Control

To maximize use of antivirals, we compare several focused strategies for antiviral use during influenza pandemics (see Figure S1 for seasonal results). Preferentially treating children ages 5–18 results in a slight reduction in attack rate when infected individuals are treated within 48 hours of symptom onset (Figure 4a). When treatment occurs within 24 hours of symptom onset, this reduction is larger. Notably, this significant reduction is achieved despite the fact that coverage levels among the entire population are lower than when antivirals are allocated randomly (compare to Figure 2b). In addition, it is key that the effect of preferentially treating children does not always increase with coverage level as all infected children are reached by 40% antiviral coverage.

**Figure 4:**
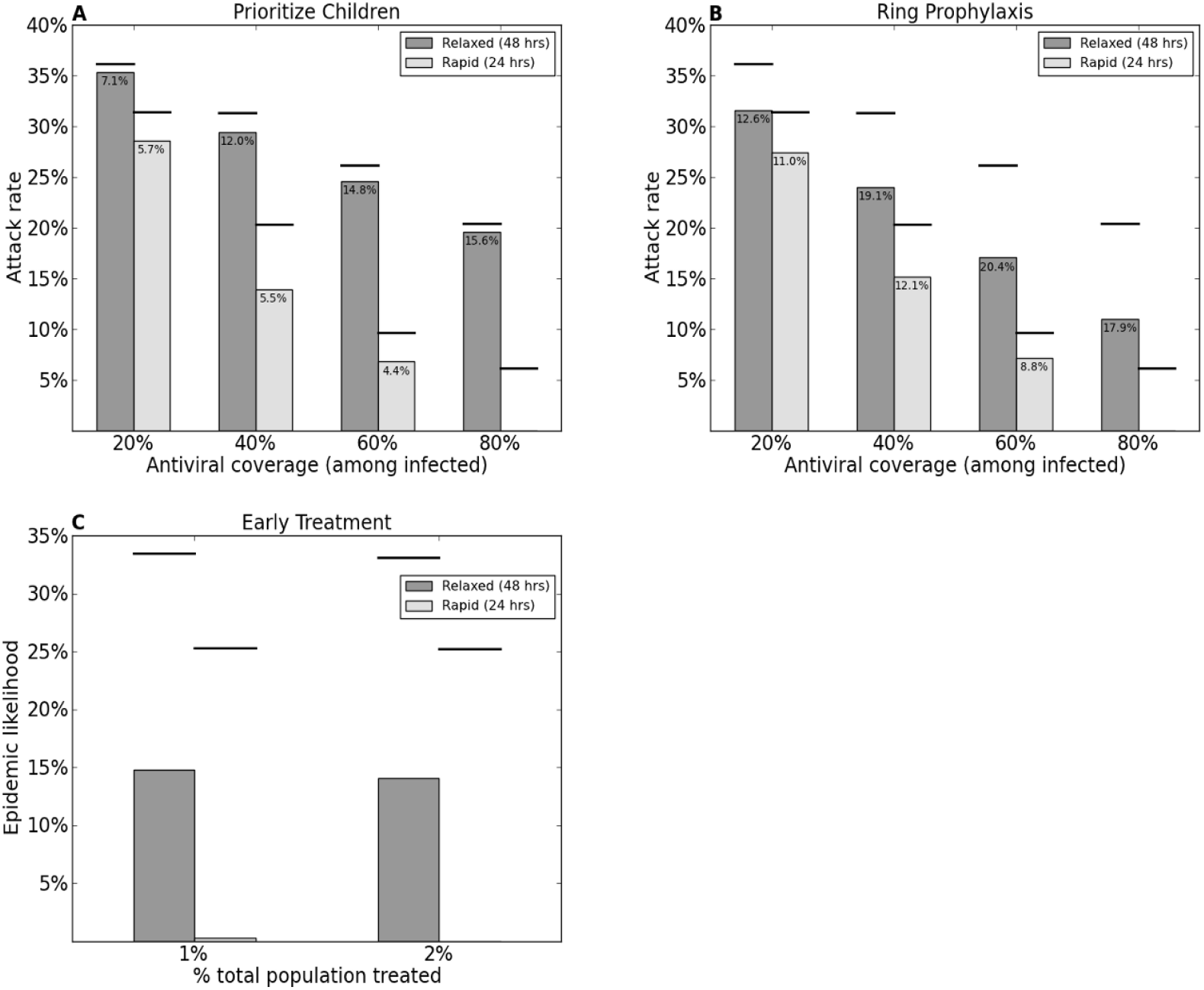
Impact of focused antiviral treatment strategies. (A,B) Attack rate (y-axis) when (A) school-age children are preferentially treated or (B) ring pro-phylaxis is used compared to random antiviral treatment (solid lines above bars) at various levels of antiviral coverage among infected individuals. Percent of entire population treated displayed within bars. (C) Epidemic likelihood (y-axis) when all infected individuals are treated until a certain percentage (x-axis) of the total population has received treatment, compared to when 30% of infected individuals are treated until the same overall coverage levels are reached (solid lines above bars). Results shown only for pandemic influenza. Simulations in which less than 5% of the population was infected are not shown. Standard errors were below 0.002 for (A-C).

When one susceptible contact of each treated infected individual is provided with prophylactic treatment, attack rate is significantly decreased in both antiviral treatment scenarios. However, while the impact of prophylaxis increases with coverage in the relaxed case, once 40% of infected individuals are treated in the rapid case, the population-level impact of prophylaxis begins to decrease. This is due to the greater efficacy of treatment in the rapid case, which prevents onward transmission regardless of prophylactic treatment. It is worth noting that the superior impact on attack rates requires higher antiviral treatment levels among the population as a whole (compare to Figure 2b).

Lastly, we consider the impact of early treatment at 1% and 2% coverage of the total population, with the aim of halting the momentum of the outbreak and preventing it from becoming a large-scale epidemic. Results are compared to epidemic likelihood when the same overall number of individuals are treated over the course of the epidemic. We note that these coverage levels are far lower than those achieved in the previous sections. The likelihood of reaching a large-scale pandemic outbreak is diminished greatly in both the relaxed and rapid cases using early treatment, and epidemics are particularly rare in the rapid case.

## 3 Discussion

Influenza antivirals are licensed for use in many countries but are not widely employed during seasonal epidemics, nor were they widely prescribed during the A(H1N1)pdm09 pandemic. In this study, our goal was to assess the potential impact of antivirals in the case of seasonal and pandemic influenza using conservative estimates of antiviral efficacy, and to determine if more focused (yet still conservative) strategies could be employed to optimize the use of antivirals. Specifically, we chose to compare the susceptibility-reducing effects of vaccines to the transmission-blocking ability of antiviral therapy.

We used a semi-empirical contact network model for an urban population to study the population-level effectiveness of antiviral treatment. Network-based models allow us to consider the individual-level and contact-level impact of reductions in susceptibility and infectivity due to interventions, and the age-structure of the population model allows for age-specific variation in efficacy, coverage, and control strategies. Our results show that antiviral treatment could significantly reduce public health burden when vaccine is either unavailable or ineffective. Compared to an influenza vaccine of moderate efficacy (approximately 65% for seasonal and 80% for pandemic), we find that antivirals administered at levels higher than 40% of all infected individuals would outperform vaccines. We note that our results represent a best-case scenario for current influenza vaccine efficacy. For seasonal influenza, vaccine efficacy varies considerably from year to year; indeed, the 2012 meta-analysis by Osterholm, et al. found that the trivalent inactivated vaccine was only significantly efficacious during eight of the twelve influenza seasons analyzed [5]. In addition, vaccine shortages occurred in over half of influenza seasons between 1999 and 2009 [6].

As an appropriate vaccine is unlikely to be available during the first wave of future influenza pandemics, intensive treatment of infected individuals with antiviral drugs could be crucial in mitigating the impact of a pandemic while a vaccine is in development. These results hold for higher pandemic transmissibility values, similar to those observed during the 1918 pandemic (Figure S2). Seasonal vaccines may be available, but, as influenza pandemics are generally caused by novel strains, such vaccines may not be efficacious enough (10–35% efficacy) against a pandemic strain to outperform realistic antiviral treatment strategies. Indeed, estimates of 2008 and 2009 seasonal vaccine efficacy against the A(H1N1)pdm09 pandemic strain vary greatly [18, 19, 20]. However, it is important to emphasize that antiviral drugs are unlikely to greatly reduce pandemic attack rates in the long run; therefore, it is still essential that an effective vaccine be developed as quickly as possible. We also emphasize that our results are based on conservative estimates (from household studies) of antiviral efficacy to reduce infectivity. Our sensitivity analyses (Figure S3) demonstrate that if reduction in infectivity is in fact higher or if future work leads to improved antivirals, the impact of antiviral therapy could greatly increase. Furthermore, we find that the comparative population-level effectiveness of antiviral treatment versus vaccination is higher when network contact heterogeneity is low (Figure S4).

When it comes to influenza vaccination, various studies have found that prioritizing certain age groups (e.g. school-age children) [21, 22, 23] or certain occupation groups (e.g. health care workers) [22, 24] is significantly preferred over random distribution. Our study finds that this is not true of antiviral distribution. Focusing on school-age children and contacts of infected individuals certainly reduces attack rates (when preferentially treating children, this effect is significantly greater with increasing antiviral efficacy, as demonstrated in Figure S5), and, when children are preferentially treated, these lowered attack rates are achieved with fewer courses of antiviral drugs. However, the additional cost of launching a focused antiviral campaign may outweigh the benefits. Antiviral treatment, in itself, is a highly-optimized strategy; by primarily treating those who are infected, it naturally captures those who are highly-connected and most likely to spread infection. Thus, it is likely that random allocation of antivirals will remain the preferred strategy during most influenza outbreaks. The timing of antiviral treatment, on the other hand, both at the individual-level (early during an individual’s infection period) and at the population-level (early during an outbreak) does have an impact worth aiming for. Efforts should be made to treat infected individuals as soon as possible, as this increases the efficacy of the drug on the individual level and prevents onward transmission, and to treat infected individuals as soon as an outbreak emerges, as this can greatly reduce the likelihood of the outbreak becoming a large-scale epidemic. These findings are in agreement with the recent results of Black et al. [16].

Our study does have some limitations. First, we do not consider explicitly the impact of antiviral therapy on severe illness or mortality. Current guidance from CDC and WHO emphasizes the importance of administering antiviral therapy to patients who are hospitalized with severe or progressive illness caused by suspected influenza and in high-risk outpatients. It is essential that antivirals continue to be used as currently recommended to treat these individuals, even when a vaccine is available. In fact, some preliminary analyses (Figure S6) indicate that focusing on those individuals at highest risk for complications and death due to influenza (e.g. the elderly and children below the age of five) could be a prudent strategy as it would result in comparable attack rates for both seasonal and pandemic influenza to those of random antiviral allocation, but would be expected to be more effective at reducing complications and death. Secondly, we do not consider the risk of antiviral resistance. Although resistance to oseltamivir and zanamivir has been limited historically [2], the 2008–2009 season brought us a strain of oseltamivir-resistant seasonal H1N1 virus that circulated globally. We plan, in future work, to consider the impact of various antiviral distribution strategies on the emergence and spread of resistance. Lastly, our seasonal influenza scenario does not incorporate pre-existing immunity, which is likely to be present in seasonal outbreaks. We expect that prior immunity would reduce the attack rates in all scenarios (naive, vaccination, antivirals) and would impact the distribution of antivirals among infected individuals, but would not change our qualitative results.

While influenza antiviral therapy is significantly effective in reducing infection burden even with random distribution, it does require active health care-seeking behavior on the part of infected individuals. A recent study by CDC found that of those with ILI during the fall wave of the A(H1N1)pdm09 pandemic, 40% of adults and 56% of children reported seeking health care for their symptoms [25]. Of the adults who sought care, 26% were diagnosed with influenza, and a further 36% were then treated with antiviral drugs [25]. Other studies have found similar but varied estimates (*∼*13%–40%) for the proportion of infected individuals who were prescribed antiviral drugs during the A(H1N1)pdm09 pandemic [11, 17]. The success of antiviral strategies thus hinges on increasing health care-seeking rates for influenza by making care accessible at locations other than hospitals and physicians’ offices, as is possible with influenza vaccination. Our sensitivity analyses (Figure S3) show that an increase in coverage levels can have a large impact on attack rates, and can compensate for low antiviral efficacy or delayed treatment initiation.

## 4 Methods

### 4.1 Population Model

We use a semi-empirical contact network model which captures the interactions that underlie respiratory disease transmission within an urban population. The model is based on demographic data from Vancouver, British Columbia, Canada [22, 26, 27]. Individuals in the network are assigned an age and age-appropriate activities (school, work, hospital, etc.). Interactions among individuals reflect household size, employment, school, and hospital data. The model population includes 10304 individuals in the following age groups: toddlers (*<* 3 years of age), preschool children (3–4), school-age children (5–18), adults (19–64), and elderly individuals (*≥* 65).

### 4.2 Epidemic and Pandemic Models

We define the transmissibility of a disease, *T*, as the average probability that an infectious individual will transmit the disease to a susceptible contact. This per contact probability of transmission summarizes the susceptibility, *σ* (e.g. immune response) and the infectivity, *ι*, (e.g. viral shedding) of individuals. The transmissibility of a given interaction is then defined as the product of infectivity of the infected individual and the susceptibility of the susceptible individuals (*T* = *ισ)*. When no intervention is implemented, *ι* = *T* and *σ* = 1 for all individuals. The transmissibility value is also linearly related to the key epidemiological parameter, *R*_0_. Transmissibility values are chosen such that seasonal influenza infects 20% of the population [1] and pandemic influenza infects 40% [28] when no control strategies are implemented. The seasonal transmissibility in our model is 0.0643 (*R*_0_ = 1.14) while the pandemic transmissibility is 0.0767 (*R*_0_ = 1.36). For comparison, basic reproduction numbers have been estimated to be about 1.2–1.4 for seasonal epidemics [1], and to be approximately 2–3 during the 1918 pandemic [29] and 1.3–1.7 during the A(H1N1)pdm09 pandemic [30].

Epidemics and pandemics are modeled using a susceptible-infected-recovered (SIR) simulation model. Beginning with an entirely susceptible population, infection is seeded at one random individual. We iteratively take each currently infected individual, infect each of its susceptible contacts with probability, *T*, and allow the infected individual to recover. 5000 such outbreaks are simulated, and an outbreak is classified as a large-scale epidemic if over 5% of the population is infected. Attack rate is defined as the proportion of the total population infected, averaged over all large-scale epidemics; and epidemic likelihood is defined as the frequency of large-scale epidemics among all outbreaks.

### 4.3 Vaccination

We model the effects of a seasonal vaccine and two pandemic vaccines, one with low coverage and high efficacy, as seen during the A(H1N1)pdm09 pandemic [31, 32], and one with high coverage and low efficacy (See Table 1). Vaccination is implemented as a reduction in susceptibility for each vaccinated individual according to age-specific efficacies of trivalent inactivated vaccine (TIV) (See Table S2 and details below).

**Vaccine Efficacy:** Most vaccine studies measure vaccine efficacy by comparing attack rates in vaccinated populations to those in unvaccinated populations, and are useful for understanding the population-level effects of vaccine. We reviewed population-level estimates for age-specific seasonal vaccine efficacy across multiple clinical trials, meta-analyses, and reviews (references in Supplement). We report the summarized results of our review in Table S3, and the vaccine efficacies used for this study in Table S2.

In our study, we incorporate vaccine efficacy at the individual-level. We define *σ* as an age-specific individual-level susceptibility. We then define *S*_v_*, the desired attack rate among individuals who have been vaccinated in the age group in question, as *S*_v_* = *S_n_*(1 − *E*), where *S_n_* is the expected attack rate among the age group when no control strategies are implemented and *E* is the population-level vaccine efficacy for the age group. We infer *σ\** for each age group by fitting *S_v_*, the true attack rate among individuals who have been vaccinated in the age group by having a reduced level of susceptibility, *σ*, to *S*_v_* using a two-type analytical percolation model [33].

For Figure 3, vaccine efficacy is gradually reduced by a factor, *r*, as,

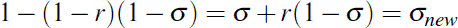

We note that a reduction in individual-level efficacy corresponds to an increase in *σ*.

**Random Vaccination:** For the seasonal scenario (Figure 2a), vaccine is distributed to 20% of the population, based on CDC data from several epidemics occurring prior to 2009 [34]. A coverage level of 30% is employed for the 2009 pandemic vaccination scenario, based on CDC coverage data for the monovalent A(H1N1)pdm09 vaccine [32], and the high coverage pandemic vaccination scenario is implemented at 40% coverage, to model increased awareness and panic which may lead to higher coverage levels in a pandemic more severe than the relatively mild A(H1N1)pdm09 pandemic. Here, all vaccines were distributed randomly, irrespective of age.

**Realistic Vaccination:** Realistic, age-specific vaccination coverage levels for both the seasonal and pandemic vaccines are derived from CDC data (Table S1; weighted averages in Table 1; references in Supplement). For the high coverage pandemic vaccine, we simply double the seasonal coverage levels for each age group. Nursing home residents are the exception, as they were already vaccinated at 90% during the realistic seasonal scenarios. Vaccine efficacy

### 4.4 Antivirals

We model the effects of antiviral treatment by reducing the infectivity of a specified proportion of infected individuals over the course of an outbreak. Individuals are selected for treatment as they became infected.

**Treatment Timing, Efficacy & Coverage:** Antiviral drugs reduce influenza transmissibility if used within 48 hours of symptom onset, and become more effective if treatment is started earlier. We model two antiviral treatment scenarios: one in which all treated individuals are treated within 48 hours (relaxed scenario), and one in which all treated individuals are treated within 24 hours (rapid scenario). We anticipate a realistic scenario to be a combination of these scenarios. Timing of treatment is modeled by altering the efficacy of antiviral drugs on individual infectivity, based on the results of household transmission studies [8, 9, 10, 11]. Thus, antiviral effectiveness was 20% for the relaxed scenario and 40% for the rapid scenario. Antiviral efficacy values are reported directly in [9]. Values were calculated from the remaining studies as *E* = 1 - (*SAR_T_/SAR_U_)*, where *E* is antiviral efficacy, and *SAR_T_* and *SAR_U_* are the reported secondary attack rates among contacts of treated and untreated individuals, respectively. If a study reported secondary attack rates separately for contacts of individuals treated within 24 hours and within 48 hours of symptom onset, efficacy values for both treatment scenarios were calculated. We report the results of our review in Table S4. Finally, averages weighted by the number of treated index cases in each study were calculated and rounded down to obtain conservative estimates.

The desired coverage is achieved by treating infected individual randomly.

**Focused Strategies:** We model three focused antiviral strategies. We model a strategy preferentially treating children (ages 5–18) by treating a certain proportion of infected children until the desired coverage level is achieved. If all infected children are treated and coverage among infected individuals remains below the desired level, antiviral drugs are distributed randomly among the remaining age groups to achieve the designated coverage level. We implement antiviral prophylaxis by treating a given proportion of infected individuals randomly, and additionally selecting randomly a single susceptible contact of each treated individual and reducing this contact’s susceptibility. We continue to randomly choose contacts of each treated individual until a susceptible contact is identified. If a treated individual has no remaining susceptible contacts, no prophylaxis is given. We assume that antiviral prophylaxis reduces susceptibility by 70% [9, 11]. Finally, we implement early antiviral treatment on the population-level by treating all infected individuals until 1%, 2%, 3%, or 5% of the entire population has received treatment. We compare this strategy to a baseline scenario in which infected individuals are treated at a realistic coverage level of 30% until the desired percentage of the entire population is treated.

## Acknowledgments

We gratefully acknowledge Lauren Ancel Meyers for use of the Vancouver urban contact network. This work was funded by a grant to Georgetown University from the Howard Hughes Medical Institute through the Precollege and Undergraduate Science Education Program; and the RAPIDD Program of the Science & Technology Directorate, Department of Homeland Security and the Fogarty International Center, National Institutes of Health.

## Disclaimer

The opinions expressed by authors contributing to this journal do not necessarily reflect the opinions of the institutions with which the authors are affiliated.

## Biographical sketches

Sarah Kramer is an undergraduate Biology of Global Health major and mathematics minor at Georgetown University. Her primary research interests include the epidemiology of and appropriate public health measures against infectious diseases.

Shweta Bansal is an Assistant Professor of Biology at Georgetown University. She is a mathematical epidemiologist and focuses on network modeling of host contact patterns and pathogen dynamics for infectious disease ecology and epidemiology.

